# Combined effects of pharmacological interventions and intermittent theta-burst stimulation on motor sequence learning

**DOI:** 10.1101/2024.07.24.604878

**Authors:** Hakjoo Kim, Paul T. Kornman, Jamie Kweon, Eric M. Wassermann, David L. Wright, Johnson Li, Joshua C. Brown

## Abstract

Drugs that modulate N-methyl-D-aspartate (NMDA) or γ-Aminobutyric acid type A (GABA_A_) receptors can shed light on their role in synaptic plasticity mechanisms underlying the effects of non-invasive brain stimulation. However, research on the combined effects of these drugs and exogenous stimulation on motor learning is limited. This study aimed to investigate the effects of pharmacological interventions combined with intermittent theta-burst stimulation (iTBS) on human motor learning. Nine right-handed healthy subjects (mean age ± SD: 31.56 ± 12.96 years; 6 females) participated in this double-blind crossover study. All participants were assigned to four drug conditions in a randomized order: (1) D-cycloserine (partial NMDA receptor agonist), (2) D-cycloserine + dextromethorphan (NMDA receptor agonist + antagonist), (3) lorazepam (GABA_A_ receptor agonist), and (4) placebo (identical microcrystalline cellulose capsule). After drug intake, participants practiced the 12-item keyboard sequential task as a baseline measure. Two hours after drug intake, iTBS was administered at the primary motor cortex. Following iTBS, the retention test was performed in the same manner as the baseline measure. Our findings revealed that lorazepam combined with iTBS impaired motor learning during the retention test. Future studies are still needed for a better understanding of the mechanisms through which TMS may influence human motor learning.

## 1. Introduction

Long-term potentiation (LTP) and long-term depression (LTD), considered fundamental mechanisms for learning and memory formation (Morris, 1989; Bliss and Collingridge, 1993; Nabavi et al., 2014), are known to be related to motor learning (Linden and Connor, 1993; Bear and Malenka, 1994; Citri and Malenka, 2008). LTP strengthens specific synapses to store information, while LTD weakens unnecessary synapses to maintain synaptic adaptability (i.e., synaptic plasticity). These processes sustain the plasticity of neural circuits, allowing adaptation to environmental changes and the learning of new information (Brown et al., 2022). Although LTP and LTD exert contrasting effects on synaptic excitability, both can be induced at the same synapse by distinct patterns of N-methyl-D-aspartate (NMDA) receptor activation (Malenka and Bear, 2004; Whalley, 2007; Lüscher & Malenka, 2012; Brown et al., 2022).

Given the crucial role of NMDA receptors in mediating both LTP and LTD, modulating these receptors presents a promising avenue for enhancing synaptic plasticity, particularly in brain disorders where plasticity is impaired. One such modulator is D-cycloserine, which has been used both as a tuberculosis antibiotic (Kuehl et al., 1955) and as a neuropsychiatric drug (Crane, 1959). D-cycloserine acts as a partial agonist at the glycine binding site of NMDA receptors in the brain (Schade and Paulus, 2016; Mataix-Cols et al., 2017). This action can enhance the function of NMDA receptors, promoting synaptic plasticity when combined membrane depolarization. Numerous studies indicate that D-cycloserine enhances neuroplasticity induced by learning and memory paradigms in both animals (Monahan et al., 1989; Flood et al., 1992; Thompson et al., 1992; Ressler et al., 2004; Davis et al., 2006) and humans (Kalisch et al., 2009; Onur et al., 2010; Kuriyama et al., 2011; Feld et al., 2013; Forsyth et al., 2015; Dempsey-Jones et al., 2021). D-cycloserine also enhances synaptic plasticity in response to 10-Hz repetitive transcranial magnetic stimulation (rTMS) (Brown et al., 2020; Kweon et al., 2022) in a manner consistent with Hebbian and homeostatic rules of LTP (Brown et al., 2021; Kweon et al., 2023; Vigne et al., 2023). D-cycloserine combined with rTMS may even be able to rescue impaired plasticity seen in depression (Cole et al., 2021, 2022).

In contrast to D-cycloserine, dextromethorphan is a non-competitive antagonist of NMDA receptors, inhibiting excessive neuronal excitation in the central nervous system (Church et al., 1985; Franklin and Murray, 1992; Netzer et al., 1993). Dextromethorphan is commonly used as a cough suppressant (Bem and Peck, 1992), but it has also been tested in the treatment of depression (Nguyen et al., 2016; Murrough et al., 2017; Majeed et al., 2021). Through NMDA receptor antagonism, dextromethorphan can inhibit the induction of LTP (Krug et al., 1993), impair learning and memory (Bane et al., 1996; Cho et al., 2006; Zhang et al., 2007; Ijomone and Biose, 2019; Bates and Trujillo, 2023), and block TMS potentiating effects (Stefan et al., 2002; Wolters et al., 2003; Wankerl et al., 2010; Weise et al., 2017).

Glutamatergic transmission and plasticity are heavily influenced by γ-Aminobutyric acid (GABA)-ergic inhibition. GABA receptors contain chloride ion channels, which hyperpolarize the membrane, preventing signal propagation and LTP (Zhu et al., 2022). As the inhibitory effect of GABA is enhanced, the excitability of the nervous system decreases, resulting in effects such as sedation, anxiolysis, anticonvulsant action, muscle relaxation, and sleep induction (Gottesmann, 2002). Lorazepam is a benzodiazepine class medication, which binds to the GABA type A (GABA_A_) receptors as a positive allosteric modulator, causing the receptor to open more frequently (Ghit et al., 2021). Wan et al. (2004) revealed that lorazepam impaired synaptic plasticity, and numerous studies suggested that lorazepam caused memory impairment (Satzger et al., 1990; Bishop and Curran, 1995; Wan et al., 2004; Pomara et al., 2015) and worsened motor performance (Bishop and Curran, 1995; Pompéia et al., 2003; Pomara et al., 2015). Another benzodiazepine, diazepam, blocked the LTP-like effects of rTMS (Heidegger et al., 2010). Interestingly, 10-Hz magnetic stimulation in mice produced both LTP-associated changes as well as a reduction in GABA_A_ receptors (Vlachos et al., 2012; Lenz et al., 2016).

We sought to study the mechanisms of intermittent theta-burst stimulation (iTBS), another excitatory protocol of TMS (Huang et al., 2005), using drugs that modulate NMDA and GABA receptors in humans. Based on observations that iTBS enhanced motor learning (Teo et al., 2011) and that NMDA receptors are both necessary (Huang et al., 2007) and sufficient to enhance rTMS-induced potentiation (Brown et al., 2020; Kweon et al., 2022), we hypothesized the following: 1) that D-cycloserine would enhance iTBS effects on motor learning performance (i.e., faster response times (RT)), 2) that this enhancement would be blocked by adding dextromethorphan, and 3) that lorazepam would impair iTBS-mediated improvements in motor learning. This study is a step towards understanding the mechanism (and augmentation potential) of iTBS on the functional expression of synaptic plasticity: learning and memory.

## 2. Materials and methods

### 2.1. Participants

A total of nine right-handed healthy subjects participated in this double-blind crossover study (mean age ± SD: 31.56 ± 12.96 years; 6 females). Exclusion criteria included individuals with any neurological disorders or those actively taking neuropsychiatric medications. Each individual was assigned to all four drug conditions in a randomized order: (1) D-cycloserine (DCS), (2) D-cycloserine and dextromethorphan (DCS + DXM), (3) lorazepam (LZP), and (4) placebo. Prior to the experiments, all participants provided written informed consent approved by the Butler Hospital Institutional Review Board. In the LZP condition, four participants experienced adverse effects from lorazepam (i.e., drowsiness). Among these four individuals, one subject was unable to complete the second session of the behavioral task (i.e., retention test) due to severe drowsiness. Consequently, the baseline data for this subject were excluded from the data analysis. Table 1 presents the demographic information of the finalized participants in each condition after the removal of missing data and outliers (see Data analysis (2.7) for the detailed method of outlier detection), resulting in loss of data for three subjects’ data from DCS condition, two from DCS+DXM, three from LZP, and one from Placebo.

**Table 1.** Demographics information in each condition. DCS: D-cycloserine; DXM: dextromethorphan; LZP: lorazepam; SD: standard deviation.

| Condition | *n* | Age (SD) | Females | Males |
| --- | --- | --- | --- | --- |
| DCS | 6 | 31.67 (12.36) | 4 | 2 |
| DCS + DXM | 7 | 31.86 (13.91) | 5 | 2 |
| LZP | 6 | 36.67 (13.22) | 4 | 2 |
| Placebo | 8 | 32.75 (13.32) | 5 | 3 |

### 2.2. Drug applications

Participants took D-cycloserine 100 mg (i.e., partial NMDA receptor agonist) in the DCS condition, D-cycloserine 100 mg and dextromethorphan 150 mg (i.e., NMDA receptor agonist + antagonist) intended to block the NMDA receptor agonism of DCS in the DCS + DXM condition, and lorazepam 2.5 mg (i.e., GABA_A_ receptor agonist) in the LZP condition, as used previously (Di Lazzaro et al., 2005, 2007; Teo et al., 2009). For the placebo condition, individuals took an identical microcrystalline cellulose capsule (Tidewater Pharmacy, Mount Pleasant, SC, United States). There was a minimum washout period of one week between each drug condition to prevent carryover effects (cf. D-cycloserine half-life: Van Berckel 1997; dextromethorphan half-life: Zhang et al., 1992; Schadel et al., 1995; lorazepam half-life: Kyriakopoulos et al., 1978; Patwardhan et al., 1980). Plasma levels of each drug were not obtained.

### 2.3. Serial reaction time task (SRTT)

In the current study, a 12-item serial reaction time task (SRTT) was used to measure the behavioral measures of synaptic plasticity through the modulation of NMDA or GABA_A_ receptors. To perform the sequence task, all participants placed the four fingers of their dominant hand (i.e., index, middle, ring, and little fingers) on the “C,” “V,” “B,” and “N” keys of a standard computer keyboard. The “C,” “V,” “B,” and “N” keys corresponded to the horizontal and parallel positions on the computer monitor where visual cues (i.e., black square) could appear, representing positions 1, 2, 3, and 4, respectively. For example, when a visual cue appeared at the far left of the screen, individuals had to press the “C” key (i.e., 1). In this study, there were a total of eight different training sequences, each as follows: Sequence #1: 2-3-1-4-3-2-4-1-3-4-2-1; Sequence #2: 1-2-4-2-3-4-3-1-3-2-1-4; Sequence #3: 3-1-2-3-1-2-3-2-1-3-4-2; Sequence #4: 4-2-3-1-4-3-2-3-2-1-3-3; Sequence #5: 1-4-2-3-2-1-4-4-3-1-2-4; Sequence #6: 2-3-1-4-1-2-4-3-1-2-1-3; Sequence #7: 4-2-3-1-3-1-4-1-3-2-3-4; Sequence #8: 3-1-3-2-4-2-1-3-1-2-4-1. Each participant practiced a different type of training sequence during each of their four visits (e.g., Sequence #1 during Visit 1, Sequence #2 during Visit 2, Sequence #3 during Visit 3, Sequence #4 during Visit 4). Participants repeated each training sequence six times consecutively and performed two 12-item random sequences before and after (i.e., R-T-T-T-T-T-T-R; R: random sequence; T: training sequence) or before the six repetitions (i.e., R-R-T-T-T-T-T-T). Each random sequence was unique, with no repetitions among them. All individuals executed a total of 96 key presses (= 12-item training sequence × 6 repetitions + 12-item random sequence × 2) during each visit.

### 2.4. Transcranial magnetic stimulation (TMS)

A PowerMAG EEG 100 TMS stimulator with a PMD70-pCool coil (2 × 70 mm) (MAG & More, Germany) was utilized for single-pulse TMS and iTBS. Single-pulse TMS was administered near the hand knob area (Yousry et al., 1997) of the primary motor cortex (M1) in the left hemisphere at the location and coil orientation experimentally determined to consistently produce the maximal motor-evoked potential (MEP) responses, termed the motor “hotspot.” After M1 determination, the resting motor threshold (rMT) was determined based on the minimum intensity that induced a peak-to-peak MEP amplitude of > 50 μV in at least five out of ten trials (Rossini et al., 2015). iTBS (burst frequency: 5 Hz, three pulses per burst at 50 Hz, a total of 600 pulses, Huang et al., 2005) was administered at the hotspot with an intensity of 80% of rMT. During the whole TMS session, the Brainsight TMS navigation system (Rogue Research, Canada) was used to keep all pulses within 0.5 mm of the labeled target accomplished through continuous manual coil positioning.

### 2.5. Electromyography (EMG)

Two disposable Ag-AgCl electromyography (EMG) electrodes (Cardinal Health, Inc., United States) were attached on the dominant hand (i.e., the active electrode at the first dorsal interosseous (FDI) muscle) to determine the rMT. The raw EMG data were amplified and filtered by the 1902 isolated pre-amplifier and Micro 1401 (Cambridge Electronic Design Limited, United Kingdom). The EMG signals were sampled and analyzed using Signal software (Cambridge Electronic Design Limited, United Kingdom). The background noise was continuously monitored.

### 2.6. Procedure

After the informed consent process, participants took the drug assigned to each group on Visits 1 through 4 (see Figure 1). Subsequently, rMT and hotspot were determined. Participants then trained on the SRTT to measure baseline behavioral data. Two hours after drug intake, iTBS was administered at M1 (i.e., hotspot of the FDI muscle). Following iTBS, individuals performed the SRTT again as a retention test and to determine whether iTBS + drug condition influenced learning. After a minimum of one week, participants were assigned to a different drug condition, and behavioral data were collected in the same manner as the first (or prior) visit. Figure 1 depicts the experimental design.

**Figure 1.**
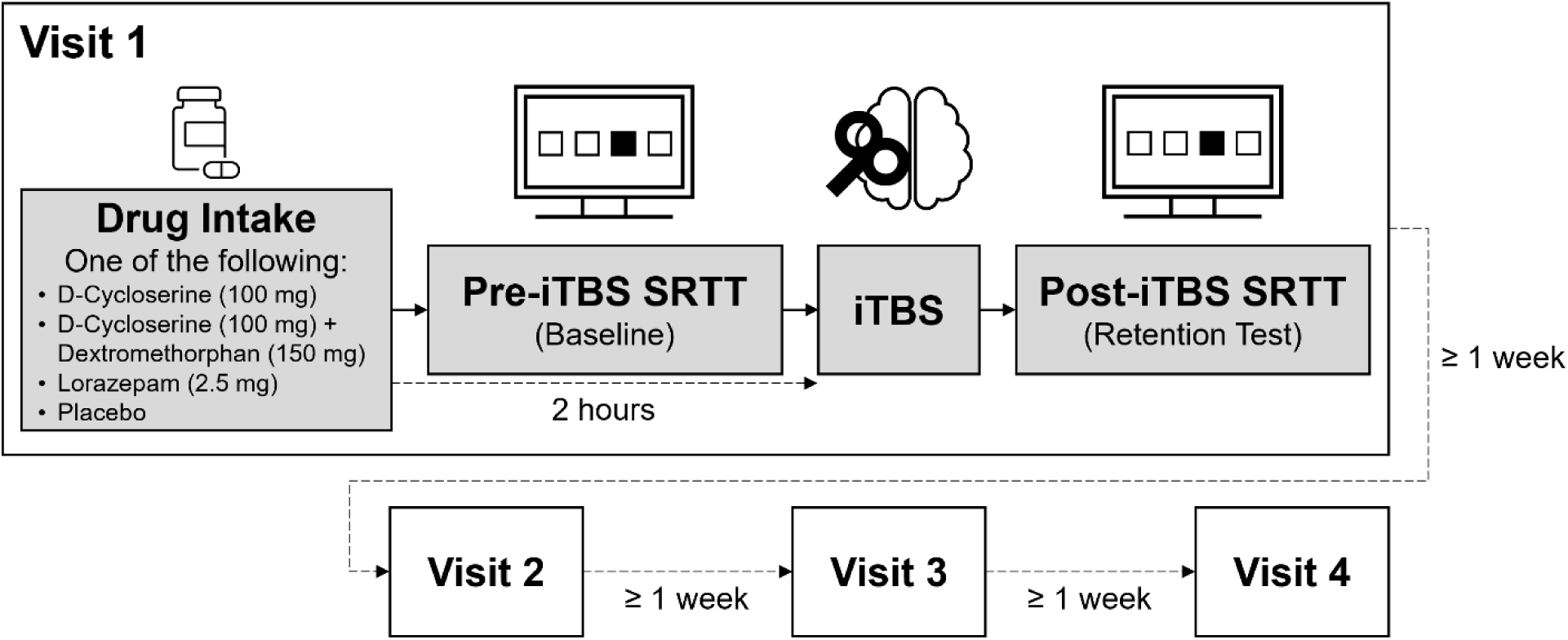
Experimental procedure. Participants trained on the SRTT after drug intake. Two hours after drug intake, they received iTBS. Following this, they performed the SRTT as a retention test and to determine whether iTBS + drug condition influenced learning. The washout period was at least one week, after which, during subsequent visits, they took a different type of drug and practiced a different sequence of the SRTT. iTBS: intermittent theta-burst stimulation; SRTT: serial reaction time task.

### 2.7. Data analysis

The primary dependent variable of the present study was the mean RT (in seconds) for a correct key press of the training sequences. The mean RT data of the random sequences were not analyzed due to the irregular ordering (i.e., R-T-T-T-T-T-T-R or R-R-T-T-T-T-T-T). The RT data of the training sequences were statistically analyzed using SPSS software (Version 28.0.0.0, IBM, United States). Outliers were detected using SPSS based on the interquartile range (IQR). Specifically, the IQR was calculated as the difference between the third quartile (Q3) and the first quartile (Q1). Data points were considered outliers if they fell below Q1 - 1.5 × IQR or above Q3 + 1.5 × IQR. Due to the small sample size, the Shapiro-Wilk test was used to determine normality. After verifying the normality of the data (*p* > .05), repeated measures analysis of variance (ANOVA) with an alpha level of 0.05 was utilized to reveal significant main effects of Trial and Drug, as well as the interaction between Drug and Trial. If a significant interaction effect was found, a simple main effects analysis was conducted, followed by post-hoc analysis using pairwise comparisons with Bonferroni adjustment. G*Power software (Version 3.1.9.7; Faul et al., 2007) was used for the post-hoc statistical power (= 1 - β (Type II error)) analysis.

## 3. Results

### 3.1. Overall performance

The 4 (Drug: DCS, DCS + DXM, LZP, Placebo) × 12 (Trial: 1 to 12) repeated measures ANOVA performed on the mean RT data revealed a significant main effect of Trial, *F*(3.726, 85.703) = 5.245 (Greenhouse-Geisser), *p* = .001, partial *η*^2^= .186, and a significant interaction effect between Drug and Trial, *F*(11.179, 85.703) = 2.935 (Greenhouse-Geisser), *p* = .002, partial *η*^2^= .277. However, the effect of Drug was not significant, *p* = .063.

Since the Drug × Trial interaction was statistically significant, a simple main effects analysis was conducted. The simple main effects of Drug on the mean RT were significant in Trials 11 (*F*(3, 23) = 5.442, *p* = .006, partial *η*^2^= .415) and 12 (*F*(3, 23) = 7.526, *p* = .001, partial *η*^2^= .495) (see Figure 2). Pairwise comparisons with Bonferroni adjustment, accounting for a total of six Drug comparisons (i.e., (1) DCS vs. DCS + DXM, (2) DCS vs. LZP, (3) DCS vs. Placebo, (4) DCS + DXM vs. LZP, (5) DCS + DXM vs. Placebo, and (6) LZP vs. Placebo), revealed the differences between Placebo and LZP in Trial 11 (*p* = .015), between DCS and LZP in Trial 11 (*p* = .009), between DCS + DXM and LZP in Trial 11 (*p* = .045), between Placebo and LZP in Trial 12 (*p* = .009), between DCS and LZP in Trial 12 (*p* = .003), and between DCS + DXM and LZP in Trial 12 (*p* = .002). In other words, a significantly slower mean RT in the LZP condition was observed starting from Trial 11. Contrary to our hypothesis about D-cycloserine, there was no significant improvement in the mean RT between the Placebo and DCS conditions across all trials.

**Figure 2.**
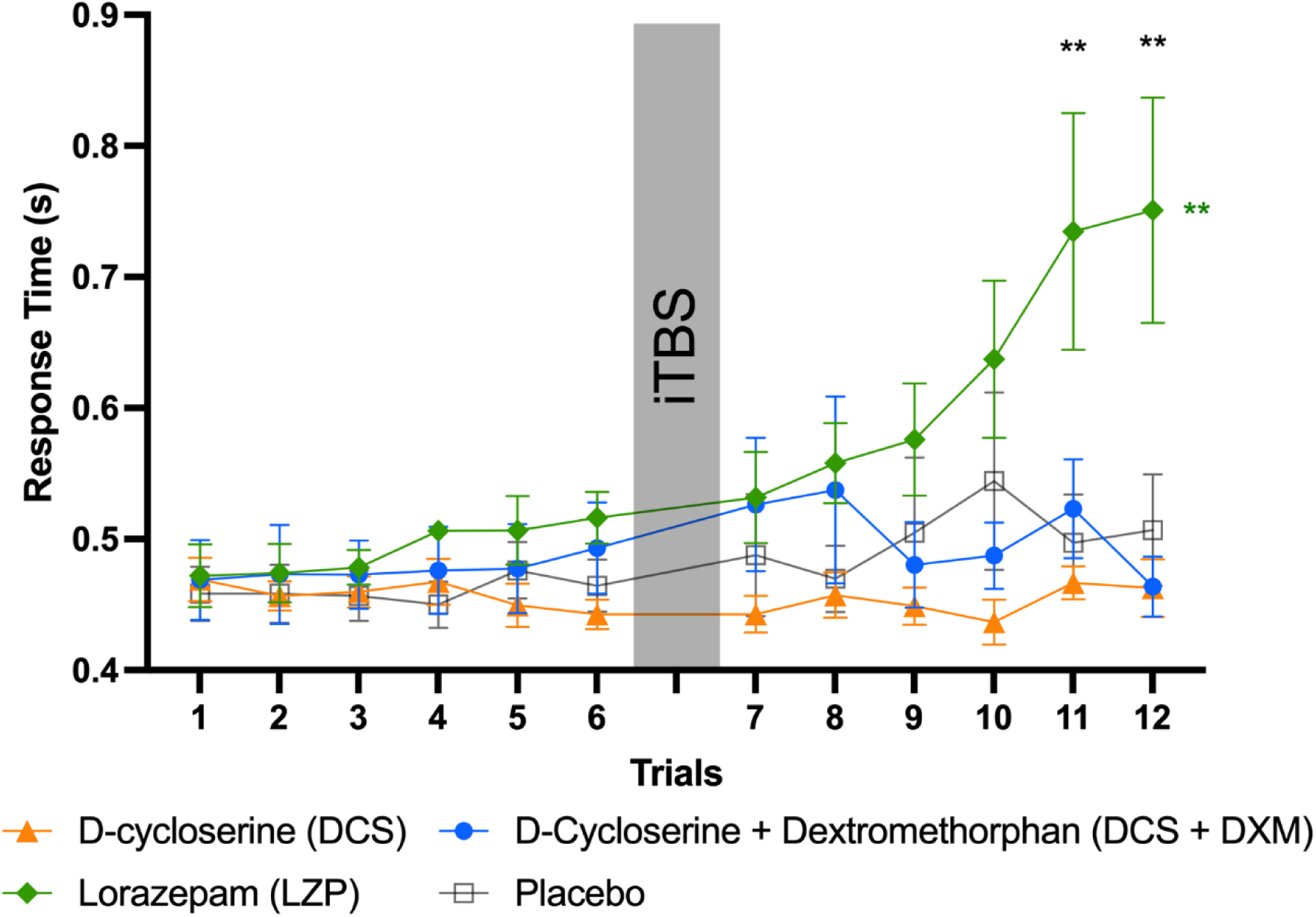
Overall task performance. Trials 1 to 6 represent the performance in the pre-iTBS training sequences, while Trials 7 to 12 represent the performance in the post-iTBS training sequences. The error bars indicate the standard errors. iTBS: intermittent theta-burst stimulation. \*\**p* < .01.

The simple main effect of Trial on the mean RT was significant in the LZP condition (*F*(11, 13) = 5.244, *p* = .003, partial *η*^2^= .816) (see Figure 2). Pairwise comparisons with Bonferroni adjustment, accounting for a total of 66 Trial comparisons (i.e., (1) Trial 1 vs. 2, (2) 1 vs. 3, (3) 1 vs. 4, …, and (66) 11 vs. 12), revealed the differences between the following trials: 1 and 11 (*p* = .001), 1 and 12 (*p* < .001), 2 and 11 (*p* = .005), 2 and 12 (*p* < .001), 3 and 11 (*p* = .003), 3 and 12 (*p* < .001), 4 and 11 (*p* = .010), 4 and 12 (*p* = .003), 5 and 11 (*p* = .023), 5 and 12 (*p* = .002), 6 and 11 (*p* = .023), 6 and 12 (*p* = .004), 7 and 11 (*p* = .011), 7 and 12 (*p* = .009), 8 and 11 (*p* = .022), 9 and 11 (*p* = .043), and 9 and 12 (*p* = .009) in the LZP condition. In other words, individuals who took lorazepam exhibited slower RTs later in the session after iTBS. This finding was consistent with our hypothesis about lorazepam.

### 3.2. Online change during pre-iTBS SRTT

Online change during pre-iTBS SRTT was defined as the difference in the mean RT between the first trial (i.e., Trial 1) and the last trial (i.e., Trial 6) in the pre-iTBS SRTT (cf. Kim et al., 2024). The 4 (Drug: DCS, DCS + DXM, LZP, Placebo) × 2 (Trial: 1, 6) repeated measures ANOVA performed on the mean RT data failed to reveal any significant main effects of Trial, *p* = .181, and Drug, *p* = .624, and a significant interaction effect between Drug and Trial, *p* = .066. Results indicate a lack of online learning over six trials regardless of drug.

### 3.3. Offline change between pre-iTBS and post-iTBS SRTT

Offline change between pre-iTBS SRTT and post-iTBS SRTT was defined as the difference in the mean RT between the last trial (i.e., Trial 6) in the pre-iTBS SRTT and the first trial (i.e., Trial 7) in the post-iTBS SRTT (cf. Robertson et al., 2004; Kim et al., 2024; Yamada et al., 2024), during which iTBS was performed. The 4 (Drug: DCS, DCS + DXM, LZP, Placebo) × 2 (Trial: 6, 7) repeated measures ANOVA performed on the mean RT data failed to reveal any significant main effects of Trial, *p* = .336, and Drug, *p* = .252, and a significant interaction effect between Drug and Trial, *p* = .934. In other words, there were no significant differences in the mean RT between Trials 6 and 7 across all drug conditions, suggesting the absence of offline learning.

### 3.4. Online change during post-iTBS SRTT

Online change during post-iTBS SRTT was defined as the difference in the mean RT between the first trial (i.e., Trial 7) and the last trial (i.e., Trial 12) in the post-iTBS SRTT (cf. Kim et al., 2024). The 4 (Drug: DCS, DCS + DXM, LZP, Placebo) × 2 (Trial: 7, 12) repeated measures ANOVA performed on the mean RT data revealed significant main effects of Trial, *F*(1, 23) = 4.598, p = .043, partial *η*^2^=.167, Drug, *F*(3, 23) = 4.023, *p* = .019, partial *η*^2^= .344, and a significant interaction effect between Drug and Trial, *F*(3, 23) = 6.542, *p* = .002, partial *η*^2^= .460.

Since the Drug × Trial interaction was statistically significant, a simple main effects analysis was conducted. The simple main effect of Drug on the mean RT was significant in Trial 12 (*F*(3, 23) = 7.526, *p* = .001, partial *η*^2^= .495). Pairwise comparisons with Bonferroni adjustment, accounting for a total of six Drug comparisons, revealed the differences between Placebo and LZP (*p* = .009), between DCS and LZP (*p* = .003), and between DCS + DXM and LZP (*p* = .002) in Trial 12.

LZP condition demonstrated significantly slower mean RT at the end of the SRTT compared to all other drug conditions. Again, contrary to our hypothesis about D-cycloserine, there was no significant improvement in the mean RT between the Placebo and DCS conditions.

The simple main effect of Trial on the mean RT was significant in the LZP condition (*F*(1, 23) = 20.809, *p* < .001, partial *η*^2^= .475, Trial 7 = 0.532 s (SD: 0.085 s), Trial 12 = 0.751 s (SD: 0.210 s)), reflecting online impairment in RT during post-iTBS SRTT as expected (see Figure 3).

**Figure 3.**
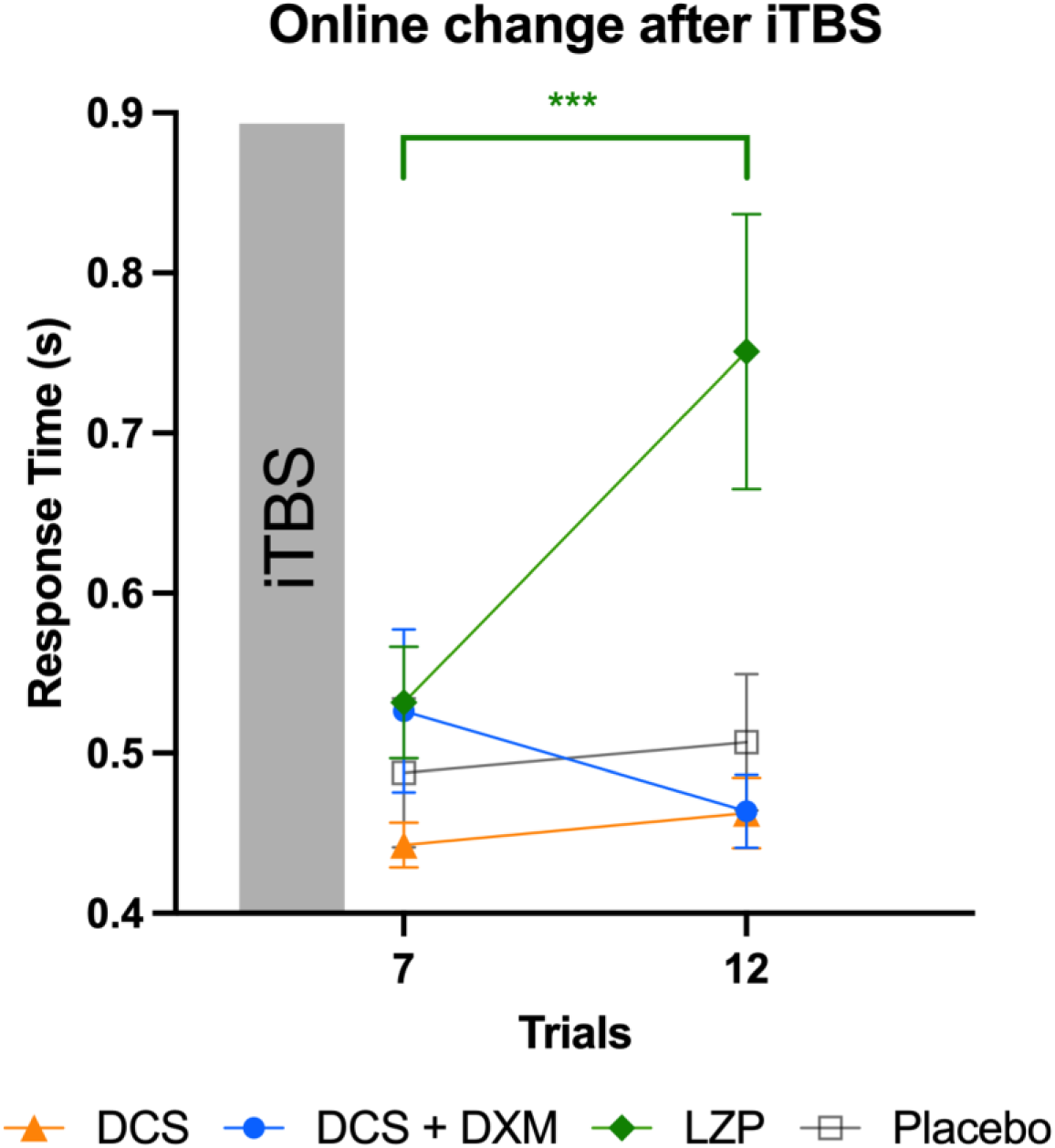
Online performance change during post-iTBS. In the LZP condition, performance worsened after practicing the task. The error bars indicate the standard errors. DCS: D-cycloserine; DXM: dextromethorphan; iTBS: intermittent theta-burst stimulation; LZP: lorazepam. *** indicates significant difference (*p* < .001) across trials in the LZP condition only.

## 4. Discussion

This study aimed to investigate the interactive effects of pharmacological interventions and M1 iTBS on human motor sequence learning. One of our hypotheses was that D-cycloserine with iTBS would produce the greatest enhancement in motor learning performance due to their synergistic effects on NMDA receptor-related synaptic plasticity (Brown et al., 2020; Kweon et al., 2022). However, we did not observe any facilitatory effects of D-cycloserine on online learning during the pre-iTBS phase, nor any combined effects of D-cycloserine and iTBS on offline or online learning during the post-iTBS phase. While previous studies have shown that D-cycloserine can enhance the excitatory effects of 10-Hz rTMS on MEP (Brown et al., 2020; Kweon et al., 2022) and could theoretically facilitate motor learning, it was not sufficient to enhance our SRTT protocol results.

A key difference between the previous studies (Brown et al., 2020; Kweon et al., 2022) and the current study was the TMS protocol. While the previous studies used 10-Hz rTMS, the present study employed the protocol of iTBS instead of 10-Hz rTMS. Although both 10-Hz rTMS and iTBS are thought to have excitatory effects, it remains unclear whether they work through identical mechanisms. Indeed, prior attempts to combine D-cycloserine with iTBS inhibited and even reversed MEP amplitude (Teo et al., 2007; Selby et al., 2019).

Baseline characteristics may influence corticomotor plasticity, which may be under- or over-represented in our small sample size. For example, in the study by Kweon et al. (2023), the group of musicians and athletes who received 10-Hz rTMS after taking D-cycloserine showed facilitated MEPs compared to the placebo with 10-Hz rTMS condition. In contrast, the non-musicians and non-athletes group showed no difference from the placebo. Another study by Vigne et al. (2023) observed that corticomotor plasticity was blunted in caffeine users, also not different from placebo.

The results of our study indicated that the combined administration of lorazepam and iTBS resulted in impairment of motor learning (see Figure 3). This is in accordance with our initial hypothesis and observations by others that demonstrated diminished motor performance due to lorazepam (Bishop and Curran, 1995; Pompéia et al., 2003; Pomara et al., 2015). Furthermore, an adverse effect common in the lorazepam condition was drowsiness. We assume that this side effect of lorazepam was caused by the decreased excitability of the nervous system as the inhibitory influence of GABA was enhanced (Gottesmann, 2002). Moreover, the GABAergic system is closely related to motor learning (Bütefisch et al., 2000; Ziemann et al., 2001; Floyer-Lea et al., 2006; Stagg et al., 2011; Kolasinski et al., 2019). For example, the GABA concentration in M1 decreased during motor sequence learning, while there were no significant changes in glutamate (Kolasinski et al., 2019). In addition, numerous studies suggested that the GABAergic inhibitory process plays a key role in motor memory stabilization (Shibata et al., 2017; Frank et al., 2022; Yamada et al., 2024). Consequently, our results demonstrated that the enhanced inhibitory effect of GABA due to lorazepam not only caused drowsiness but also impaired their performance.

Despite these findings, this study has several limitations that need to be acknowledged. One limitation is the small sample size, which may have prevented us from detecting the pharmacological effect of D-cycloserine on motor learning. According to the post-hoc power analysis, with a partial eta-squared effect size of .277, 34 subjects per group were required to achieve a power of 0.81 for the interaction effect in the overall performance analysis. In the case of the analysis of the online change during the pre-iTBS SRTT, nine subjects per group were needed to achieve a power of 0.82. For the offline change analysis, 150 subjects per group were required to achieve a power of 0.80. However, it is important to note that the result of the power analysis for the offline change may suggest that the actual effect is either very small or nonexistent. For the analysis of the online change during the post-iTBS SRTT, a power of 0.88 could be achieved with only five subjects per group due to the large effect size of lorazepam.

Another consideration that may explain null D-cycloserine results, as shown in Figure 2, six repetitions of the 12-item sequence were not sufficient to observe a clear learning curve. For example, other SRTT studies that use 12-item sequences had 15 or 25 repetitions (i.e., 180 or 300 key presses) for training and 15 repetitions (i.e., 180 key presses) for test block(s) (Cohen et al., 2005; Brown and Robertson, 2007; Cohen and Robertson, 2007). Future research should address these limitations by ensuring a sufficient amount of motor training to induce neural plasticity and observable learning. This will provide a clearer understanding of the mechanisms through which TMS influences human motor learning and underscore their potential clinical implications, such as in depression treatment.

## 6. Author Contributions

HK: Data curation, Formal analysis, Validation, Visualization, Writing – original draft, Writing – review & editing; PTK: Data curation, Investigation, Software, Writing – review & editing; JK: Data curation, Investigation, Writing – review & editing; EMW: Funding acquisition, Software, Writing – review & editing; DLW: Formal analysis, Validation, Writing – original draft, Writing – review & editing; JL: Formal analysis, Validation, Writing – review & editing; JCB: Conceptualization, Data curation, Formal analysis, Funding acquisition, Methodology, Project administration, Resources, Supervision, Validation, Writing – original draft, Writing – review & editing.

